# Interaction modulation through arrays of clustered methyl-arginine protein modifications

**DOI:** 10.1101/289041

**Authors:** Jonathan Woodsmith, Victoria Casado-Medrano, Nouhad Benlasfer, Rebecca L. Eccles, Saskia Hutten, Christian L. Heine, Verena Thormann, Claudia Abou-Ajram, Oliver Rocks, Dorothee Dormann, Ulrich Stelzl

## Abstract

Systematic analysis of human arginine methylation identifies two distinct signaling modes; either isolated modifications akin to canonical PTM regulation, or clustered arrays within disordered protein sequence. Hundreds of proteins contain these methyl-arginine arrays and are more prone to accumulate mutations and more tightly expression-regulated than dispersed methylation targets. Arginines within an array in the highly methylated RNA binding protein SYNCRIP were experimentally shown to function in concert providing a tunable protein interaction interface. Quantitative immuno-precipitation assays defined two distinct cumulative binding mechanisms operating across 18 proximal arginine-glycine (RG) motifs in SYNCRIP. Functional binding to the methyl-transferase PRMT1 was promoted by continual arginine stretches while interaction with the methyl-binding protein SMN1 was arginine content dependent irrespective of linear position within the unstructured region. This study highlights how highly-repetitive modifiable amino acid arrays in low structural complexity regions can provide regulatory platforms, with SYNCRIP as an extreme example how arginine methylation leverages these disordered sequences to mediate cellular interactions.

## Introduction

Protein post translational modifications (PTMs) are known to regulate a vast array of cellular processes governing all facets of human biology. A general three tier system of PTM addition or removal enzymes (writers and erasers) and PTM binding proteins (readers) is utilized in a wide variety of differing flavors to vastly increase the functional complexity of the human (Chen et al. 2011; Khoury et al. 2011). Substrates can be targeted by single or multiple modifications at any given time, leading to alterations in expression, localization, activity or binding partner profiles (Woodsmith and Stelzl 2014). The collection of tens of thousands of annotated sites has aided computational systematic analysis into both their evolution and interplay with one another (Beltrao et al. 2012; Minguez et al. 2012; Woodsmith et al. 2013). In particular, recorded protein arginine methylation events increased in recent years facilitating their systematic study (Guo et al. 2014; Geoghegan et al. 2015; Sylvestersen et al. 2014; Bremang et al. 2013; Larsen et al. 2016).

While issues remain with robust identification of methylation sites (Hart-Smith et al. 2016), both high-throughput dataset collections and small scale studies (for an extensive review see (Biggar and Li 2015)) highlight that arginine mono- and di-methylation impact a wide range of biological processes. Indeed, a recent large scale study identified that at least 7% of arginines in the expressed proteome are mono-methylated (Larsen et al. 2016). Comprehensive protein methylation specific interaction networks (Weimann et al. 2013) and methyltransferase knock out studies in cell culture (Shishkova et al. 2017) are beginning to define a wide array of molecular targets to support genetic studies showing the broad impact of the nine identified arginine methyl-transferase enzymes (PRMTs) *in vivo*. PRMT1 and PRMT5 have been shown to be of critical importance, displaying embryonic lethality upon knock out (Pawlak et al. 2000; Tee et al. 2010), with the majority of other PRMTs showing different forms of developmental or cellular defects (reviewed in detail in (Blanc and Richard 2017)). Furthermore PRMTs are well-documented to be dysregulated in cancer, with over-expression of PRMT1, CARM1 (PRMT4) and PRMT5 observed in several studies (Yang and Bedford 2013).

On a mechanistic level, the relationship between reported methyl-arginine sites and their cognate reader and writer proteins has been previously studied largely using short synthesized peptides *in vitro.* For example, PRMT1 and PRMT6 have been shown to prefer, but are not limited to, arginine-glycine motifs (RG/RGG motifs (Thandapani et al. 2013), referred to as RG motifs from here onwards) while CARM1 preferentially targets proline flanked arginines (Osborne et al. 2007; Kölbel et al. 2009; Gui et al. 2013). The methyl-arginine binding TUDOR domain has been annotated across 15 proteins to date, with the isolated TUDOR domain in key splicing regulator SMN1 showing a binding preference for methylated RG motif containing peptides. Furthermore, isolated TUDOR domains bind peptides with multiple modifications with a higher affinity than those with only a single methyl-arginine (Tripsianes et al. 2011; Liu et al. 2012). Indeed, many proteins have now been defined with multiple arginine methylation sites (Larsen et al. 2016), yet the potential interplay between modifications across full length sequences remains poorly studied. Furthermore, how any cooperation between modified residues mechanistically mediates specific binding preferences in the context of a writer-substrate-reader relationship in human cells is as yet unclear.

PTMs have been shown to cluster within intrinsically disordered regions of proteins, a prevalent feature throughout the proteome (Woodsmith et al. 2013). Select few of these regions have been extensively studied and experimental insight into the regulation of the majority of these unstructured regions is limited. Indeed, while subsequent bioinformatic studies have improved ways in which to identify functional PTM clusters through integration of distinct data types (Dewhurst et al. 2015), dissecting them mechanistically has proved a major challenge. *In vitro* peptide studies have provided insight into biophysical binding properties of short modified sequences but cannot address the full complexity of the long sequences identified *in vivo*. As the long intrinsically disordered protein sequences that harbor these regions lay outside of the classical structure-function paradigm, novel approaches to understanding their regulation in a cellular context are required. Furthermore, given the vast array of human proteins that contain modified disordered regions are also implicated in neurodegenerative disorders and cancer, understanding how such large regions of low structural complexity are utilized as regulatory elements is paramount to a better understanding of human cell biology (Babu 2016).

Here we highlight that arginine methylation can be broadly separated into two classes based on clustering prevalence, either isolated or within modification arrays. The existence of two distinct classes in methylated residues is supported by differences in structural context, mutational signatures and expression analysis of target proteins. We then experimentally dissected in detail the functional requirement of a highly methylated unstructured region in the heterogeneous nuclear ribonuclear protein (hnRNP) SYNCRIP. To achieve a comprehensive overview of the entire disordered region, we took a genetic approach to define the binding preferences of a stretch of 19 arginines in the C-terminal SYNCRIP tail using a panel of 37 full length mutants in quantitative immuno-precipitation experiments. To define both the unmodified and modified states of the arginine array, we leveraged the methyl-transferase PRMT1 and the methyl-binding protein SMN1 as functional readouts for arginine and methyl-arginine respectively. Remarkably, the exact same protein sequence can mediate distinct cumulative binding mechanisms in the modified and un-modified states. While both interactors increased binding concomitantly with arginine content, unmodified arginines are preferred in continual stretches in direct contrast to their modified counterparts that function in concert irrespective of their position within the structurally disordered array.

This study reveals how extensive RG repeats within low structural complexity regions can generate cumulative binding mechanisms and furthermore how extensive post translational modifications allow for a second, distinct recognition mode in a single repeat region.

## Results

### Systematic characterization of methyl-arginine array containing proteins

To investigate systematic trends of protein methylation, we initially obtained a list of all arginine and lysine methylation sites available through PhosphoSitePlus (downloaded from PhosphoSitePlus.org June. 2017). These PTMs were then mapped to unique Refseq identifiers to give 9339 arginine modifications and 4555 lysine modifications (Supplementary Table 1). We and others have previously shown that PTMs can cluster across linear protein sequences (Beltrao et al. 2012; Woodsmith et al. 2013), a finding that has been extended to 3D protein structures (Dewhurst et al. 2015). While protein structures provide a more detailed viewpoint from which to study PTM distributions, they are inherently biased against unstructured regions as well as limited in number, and as such would impose a large constraint on the PTM dataset. We therefore performed a sliding window analysis that counted the number of modified residues in stretches of 20 amino acids across a linear protein sequence (Material and Methods). The proportion of total lysine methylation that accumulates in short sequence stretches is consistently lower than that of arginine methylation across multiple modification cut-offs (Figure 1A). To systematically characterize these methylated arginine clusters, we initially investigated their sequence context. As approximately 31% of arginine methylation sites from HEK293T cells were recently shown to be contained within RG motifs (Larsen et al. 2016), we analyzed the propensity of this motif within these clustered sites. While more dispersed arginine methylation sites (1 or 2 methyl-Rs / 20 amino acid window) recapitulate this approximate 30% RG motif content, increasing densities of methylation sites correlate with a noted increase in RG motifs, up to 54% for ≥4 methylation sites per window (Figure 1B). These clustered, RG-motif driven methylation sites also correlate with a large shift towards structurally disordered regions in comparison to isolated methyl-Rs (Figure 1C).

**Figure 1.**
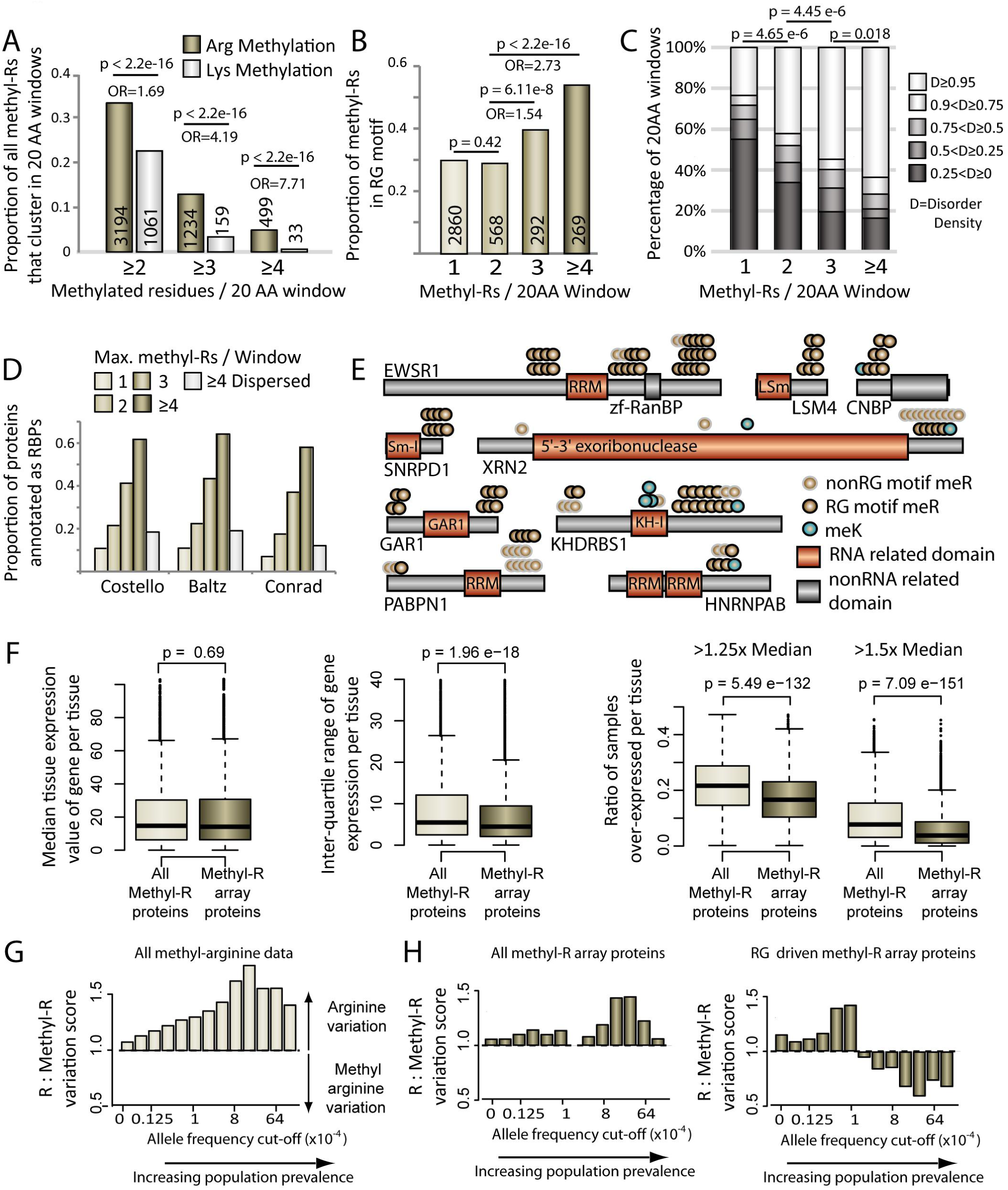
Methyl-arginine clusters show distinct sequence, expression and mutational signatures. A Bar graph showing the percentage of methyl-arginine or methyl-lysine that clusters within 20 amino acid windows. B Proportion of methyl-arginines that are contained within an RG motif in each class of 20 amino acid window. P Values and Odds Ratio (OR) for A and B calculated using Fishers Exact Test, numbers within the bars are the total methylation sites used to calculate each bar. C Bar graph showing the disorder distribution in 20 amino acid windows defined by the number of methyl-argnines they contain. P values calulated using two sided Mann-Whitney-Wicoxon test. D Bar graph showing the ratio of each protein class that are annotated as RBPs in each of the 3 named studies. Dispered methyl-arginine targets have 4 methylation sites but no clustering propensity. E Schematic diagrams of clustered methylation target proteins. F Median expression, inter-quartile range and overexpression analysis (Left pair 1.25x median cut-off, right pair 1.Sx median cut-off) of clustered methyl-arginine proteins and an equal size randomly sampled comparison group from all other methylated proteins. P values calulated using two sided Mann-Whitney-Wicoxon test. G Barchart showing the ratio of arginines that have an annotated mis-sense variant in comparison to the ratio of methyl-arginines that have an annotated mis-sense variant at differing allele frequency cut-offs. H As in G but only using arginines present in proteins targetted by clustered methyl-arginine windows (All data left, RG driven windows right).

Arginine methylated proteins have been shown to be involved in multiple facets of RNA processing and binding, for example proteins containing RRM and RH RNA binding domains are preferentially modified (Larsen et al. 2016). We therefore examined the prevalence of methyl-arginine clusters across three large scale RNA binding protein (RBP) PAR-Clip studies, which have defined the RNA binding protein repertoire (Castello et al. 2012; Baltz et al. 2012; Conrad et al. 2016). We classified protein methylation targets based upon maximum methyl-R clustering and observed a sharp increase in the fraction of targeted proteins annotated as RBPs with increasing modification density (Figure 1D). This is likely a function of clustered modifications, as proteins targeted by many, yet dispersed, arginine methylation events have a vastly reduced RBP annotation ratio (Figure 1D).

We next sought to define methyl-R clusters in full length protein sequences. We therefore scanned across each modified protein sequence to further identify proteins that contained multiple or extended arrays over and above a 20 amino acid window (Material and Methods). Using a cut-off of ≥3 proximal modifications we systemically defined 313 methyl-R arrays distributed across 273 proteins, containing a total of 1,600 arginine methylation sites (Supplementary Table 2). These arrays are distributed over a broad size range up to 182 amino acids in length, with 92 proteins having a methyl-R array longer than 25 residues (Supplementary Figure 1A). Several proteins contain multiple methyl-R arrays, such as the RNA processing proteins EWSR1 and GAR1 (Figure 1E). While RG motifs are highly prevalent in many arrays, motif analysis of non-RG driven methylation arrays suggest that CARM1 may also mediate modification clustering (Supplementary Figure 1B).

We next sought to further characterize these targets of clustered arginine methylation through comparison with their non-clustered methyl-R counterparts. As highlighted above, methyl-R arrays largely appear in regions of low structural complexity away from classical function protein domains (Figure 1C). Proteins containing these disordered regions have been shown to be under tight expression regulation in lower eukaryotes (Vavouri et al. 2009; Gsponer et al. 2008). We therefore utilized the recent GTEx gene expression dataset (Battle et al. 2017) to observe whether this trend holds for human genes targeted by clustered arginine methylation. In the GTEx dataset, each gene is associated with multiple individual samples per tissue, allowing characterization of expression variance across individuals within multiple distinct cellular environments. We first established an analytical framework to control for overall expression patterns of targeted proteins. We then characterized the median expression values of clustered methyl-R targets across 51 distinct tissues (Clustered Methyl-R Proteins Figure 1F, Material and Methods). For comparison we sampled the same number of genes from the non-clustered methyl-R target genes, using a randomization protocol that generated a statistically indistinguishable control dataset (All Methyl-R proteins, Figure 1F). When comparing the interquartile range of these two groups, we observed overall that proteins containing methyl-R arrays have more tightly controlled gene expression variance (middle panel, Figure 1F). Furthermore, the ratio of samples that show overexpression for a given gene is lower for the methyl-R array containing proteins (Cut-off 1.25x & 1.5x median expression value, rightmost panel Figure 1F). This analysis suggests that the two classes of methylation target proteins discovered based on PTM clustering also show a distinguished gene regulatory signature, with proteins harbouring methyl-R arrays under more tight expression regulation across tissues.

Finally, we turned to examine patterns of genetic variation on both classes of arginine methylation. Large scale genome and exome sequencing events have recorded the population prevalence (allele frequency) of millions of genetic variants in healthy individuals (Lek et al. 2016). These allele frequencies can act as proxies for the importance of specific amino acid residues affecting critical protein functions (Woodsmith and Stelzl 2017); in general, critical residues should seldom be targets of mis-sense mutation in healthy individuals. Here, non-methylated arginines in targeted proteins act as control for amino-acid and gene specific mutation rates. When comparing the proportion of mutated non-modified arginines to mutated methyl-arginines across the entire methylation dataset, we observe an increasing ratio indicating a relative increase of mutated non-modified arginines at increasing population prevalence (left panel, Figure 1G). This is to be expected, as the exact identity of individual post translationally modified residues will generally be more critical than their non-modified counterparts. Interestingly, when repeating this analysis for arginines contained within methyl-R arrays, this trend is substantially reduced (left panel, Figure 1H). Furthermore, when we look at those methyl-R arrays driven by RG motifs, the ratio of mutated modified arginines is actually higher than its non-modified counterpart at higher allele frequencies (right panel, Figure 1H). This analysis indicates that arginine residues targeted by methylation present in arrays are more variable in comparison to the bulk of methylated arginines outside of methyl-R arrays regions. As such, in the context of arginine methylation, the exact amino acid identity at any position where an arginine is present in these arrays is likely less critical than of more isolated methyl-arginines counterparts.

In summary, based on sequence, regulatory and genetic signatures this systematic *in silico* analysis provides evidence that protein arginine methylation occurs in two classes; methylation events that act in structurally complex regions but in relative isolation to one another, and arrays of arginine methylation where modifications may act in concert to regulate otherwise structurally less defined, low information protein sequences. To understand mechanistically how such extensive stretches of modifications can function in the cell, we sought to experimentally characterize a methyl-R array containing protein.

### Identification of methyl-arginine binding proteins for highly methylated hnRNP SYNCRIP

Utilizing short chemically modified peptides *in vitro* has shown that clusters of up to 4 methylated arginine residues distributed across 20 amino acids can markedly increase methyl binding domain interaction affinity (Tripsianes et al. 2011). Yet it is currently completely unclear whether low structural complexity regions utilize extensive methyl-R arrays stretching over dozens of amino acids for multiple independent regulatory events or whether they cumulatively combine to increase the regulatory capacity of the entire region. Not only are many of these large methyl-R arrays unamenable to *in vitro* peptide studies, it is of considerable interest how large low structural complexity regions receive and transmit information in the absence of defined structure (Babu 2016). We therefore sought a highly methylated protein to be able to characterize these extensive disordered regions in a larger context.

Previously we identified candidate methyl-transferases for a large panel of target proteins using Y2H-Seq (Weimann et al. 2013). We cross referenced proteins with the highest methylated arginine density with the arginine methyl transferase interaction results and identified the hnRNP SYNCRIP (HNRNPQ), a robust PRMT1 interactor, in the intersection for detailed hypothesis driven investigation (Supplementary Figure 1B). SYNCRIP has a total of 18 putative RG methylation target motifs spread across 106 amino acids within its disordered C terminal tail (Figure 2A & B, plus one R followed by an A). 15 of the arginines have been shown previously to be methylated spanning the entire length of its C terminal tail both *in vitro* and *in vivo* by PRMT1, 7 of which by independent studies (to date 8 arginines observed in both mono- and di-methylated state, 6 in the mono-methylated state only, and 1 on in the di-methylated state (Hornbeck et al. 2015; Larsen et al. 2016; Weimann et al. 2013). Five arginines within the array have been shown to be mutated in healthy individuals (Figure 2A, lower panel & Supplementary Table 3), and SYNCRIP shows very tight expression regulation (Figure 2C), identifying it as a true representation of the bioinformatic trends observed above.

**Figure 2.**
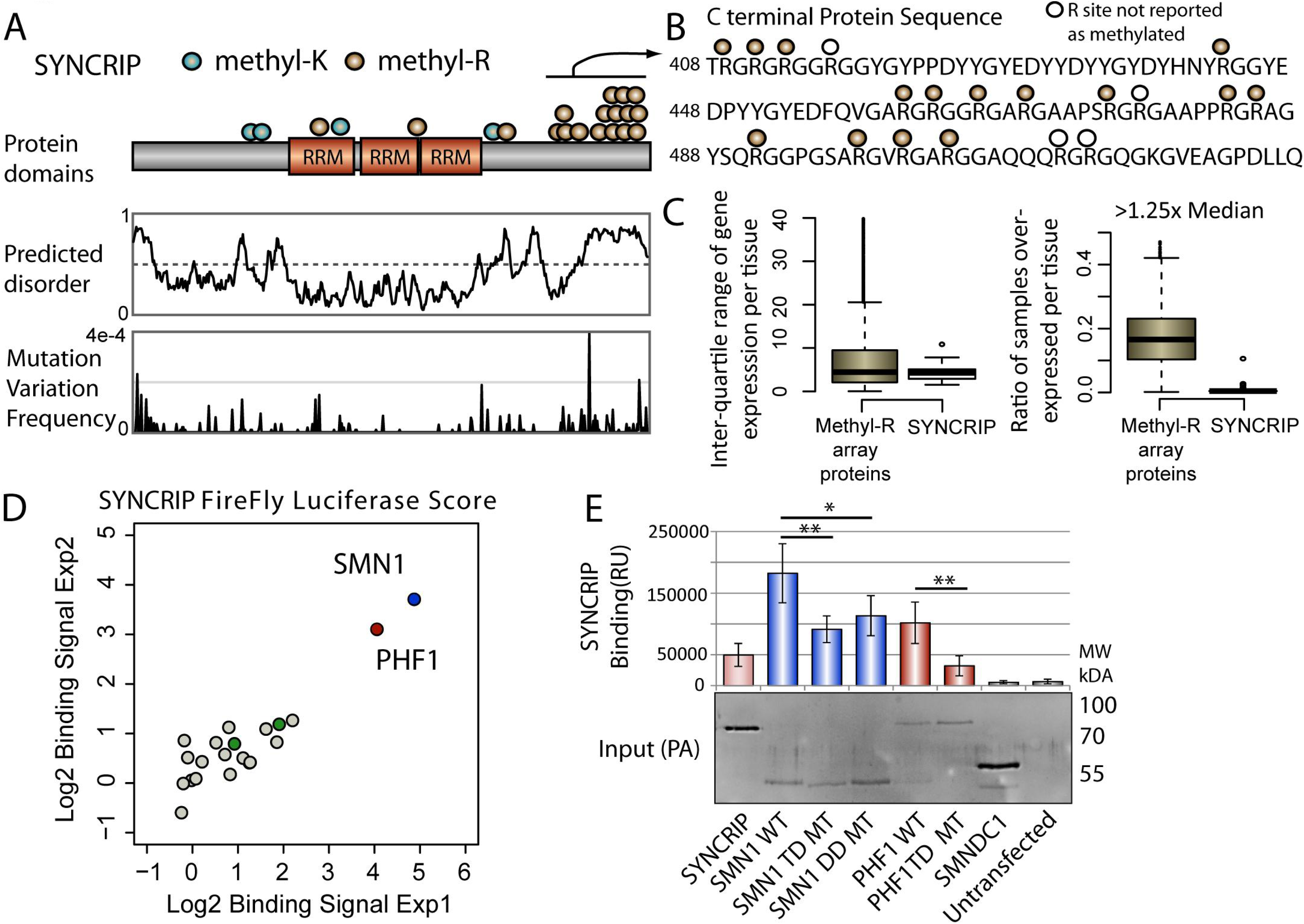
Identification of MeBPs that interact with SYNCRIP. A Schematic representation of methylation sites annotated on the hnRNP SYNCRIP. Middle panel: Predicted protein disorder of SYNCRIP. Lower Panel: Population Max frequencies of genetic variation across SYNCRIP taken from the Gnomad dataset. For ease of representation y-axis is truncated to 4e-4. B Amino acid sequence and arginine modifications in the C terminal tail of SYNCRIP. C SYNCRIP GTEx expression data: lnter-quatile range values (left panel) and overexression ratios (right panel), for comparison with data obtained for all clustered methyl-R target proteins. D LUMIER experiments showing SMNl and PHFl binding signals separate from the majority distribution of other tested MeBPs. Green dots represent non-binding TUDOR domain containing MeBPs (SMNDCl and PHF19).**E** LUMIER experiment testing SMNl and PHFl mutants against wild type SYNCRIP. TD= TUDOR domain, DD= Dimerisation domain. Input western blot to show expression of PA tagged constructs. Error bars represent highest and lowest observed values, Two sided Mann-Whitney-Wilcoxon Test used to determine statistical significance.*= P<0.05,**= P<0.01.

Based on previous structural and biochemical studies of methylated arginines present in RG type repeats, we hypothesized the C terminal tail of SYNCRIP was required for binding to one or more methyl binding domain containing proteins (MeBPs). We screened full length protein-A tagged SYNCRIP against a panel of 21 luciferase tagged putative or bona-fide MeBPs in a high throughput immunoprecipitation LUMIER-type assay (Hegele et al. 2012), allowing a quantitative readout of multiple protein-protein interactions in a 96 well format (see Material and methods). The vast majority of MeBPs showed only low signal in the LUMIER experiment, representing background binding in the assay (Figure 2D). While two of the four TUDOR domain containing proteins tested showed no interaction signal (green dots Figure 2D, SPF30 (SMNDC1) and PHF19), SMN1 and PHF1 both showed high interaction readout clearly distinct from the background distribution and were verified across repeat assays (Figure 2D). SMN1 has been previously observed to interact with multiple unrelated methylated arginine peptide sequences (Liu et al. 2012; Friesen et al. 2001; Tripsianes et al. 2011) and reported to interact with full length SYNCRIP (Rossoll et al. 2002), but any methylation dependency of the SYNCRIP-SMN1 interaction is unclear. The PHF1 TUDOR domain has been structurally characterized in complex with a histone 3 derived methyl-lysine peptide (Musselman et al. 2012). As SYNCRIP has been reported to be both lysine and arginine methylated (Figure 2A), we tested the methylation dependency of both interactors by mutating residues critical for methyl binding in the β-barrel TUDOR structure of each MeBP (TD mutants, Supplementary Figure 2A and B (Tripsianes et al. 2011; Musselman et al. 2012)). We tested these TD mutants using the LUMIER approach alongside a disease associated mutant perturbing SMN1 dimerisation that is critical for function (DD mutant, (Burghes and Beattie 2009)). Mutations in the β-barrel structure markedly reduced the SYNCRIP interaction signal without affecting the expression of either protein, suggesting these interactions are methylation dependent (TD mutants, Figure 2E). While SYNCRIP shows no self-interaction in this assay, robust SMN1 homo-oligomerization is required for a wild type SYNCRIP binding signal (DD mutant Figure 2E, and Supplementary Figure 2C). This is in line with previous literature suggesting functional hnRNP particles are disrupted by mutations in the SMN1 oligomerization domain (Burghes and Beattie 2009). In the SMN1 binding assay the TD mutant also showed a reduced dimerization signal (Supplementary Figure 2), therefore we looked for further evidence to support the methylation dependency of the interaction. Furthermore, as this study focused on the function of methyl-R arrays in protein sequences, the likely methyl-lysine dependent SYNCRIP-PHF1 interaction was not pursued further.

### SYNCRIP arginine methylation function in cell culture

To characterize the function of the SYNCRIP methyl-R array in cells, we created HEK293T cells stably expressing HA-STREP tagged wild type and lysine to arginine mutant SYNCRIP. Here we generated stable cells lines expressing arginine to lysine mutated SYNCRIP with either a small or intermediate number of the original arginines remaining in the C-terminal tail (6 and 14 arginines respectively. 6Rs mutant, remaining Rs: R409, R411, R413, R416, R475, R477. 14 Rs mutant, 5 mutants; R443K, R475K, R477K, R511K, R513K). We then immuno-precipitated exogenously expressed wild type and mutant SYNCRIP under basal conditions in HEK293T cells to assay its methylation status and endogenous SMN1 binding. Immuno-precipitated wild type SYNCRIP showed a strong signal with the pan-methylated-arginine antibody under basal conditions in HEK293T cells, a signal that was markedly reduced by the chemical methylation inhibitor Adox (Figure 3A). In support of the methylation dependent nature of the interaction, precipitated endogenous SMN1 signal was abolished in the presence of the inhibitor (Figure 3A). This pharmacologically inhibited methylation signal was mimicked by reducing the number of arginines present in the C terminal tail of SYNCRIP (two rightmost lanes Figure 3A). Methylation was undetectable above background levels on the 6R mutant, yet was partially rescued in a mutant containing 14 arginines. In agreement with the pull down of wild type SYNCRIP in the presence of Adox, mutant SYNCRIP containing only 6 arginines in the C-terminal tail showed no SMN1 binding above background levels, while the interaction was partially rescued in the mutant containing 14 C-terminal tail arginines. This experiment importantly shows that SYNCRIP can be methylated by endogenous PRMTs under basal conditions and is subsequently bound by endogenous SMN1 in a methylation dependent manner in mammalian cells.

**Figure 3.**
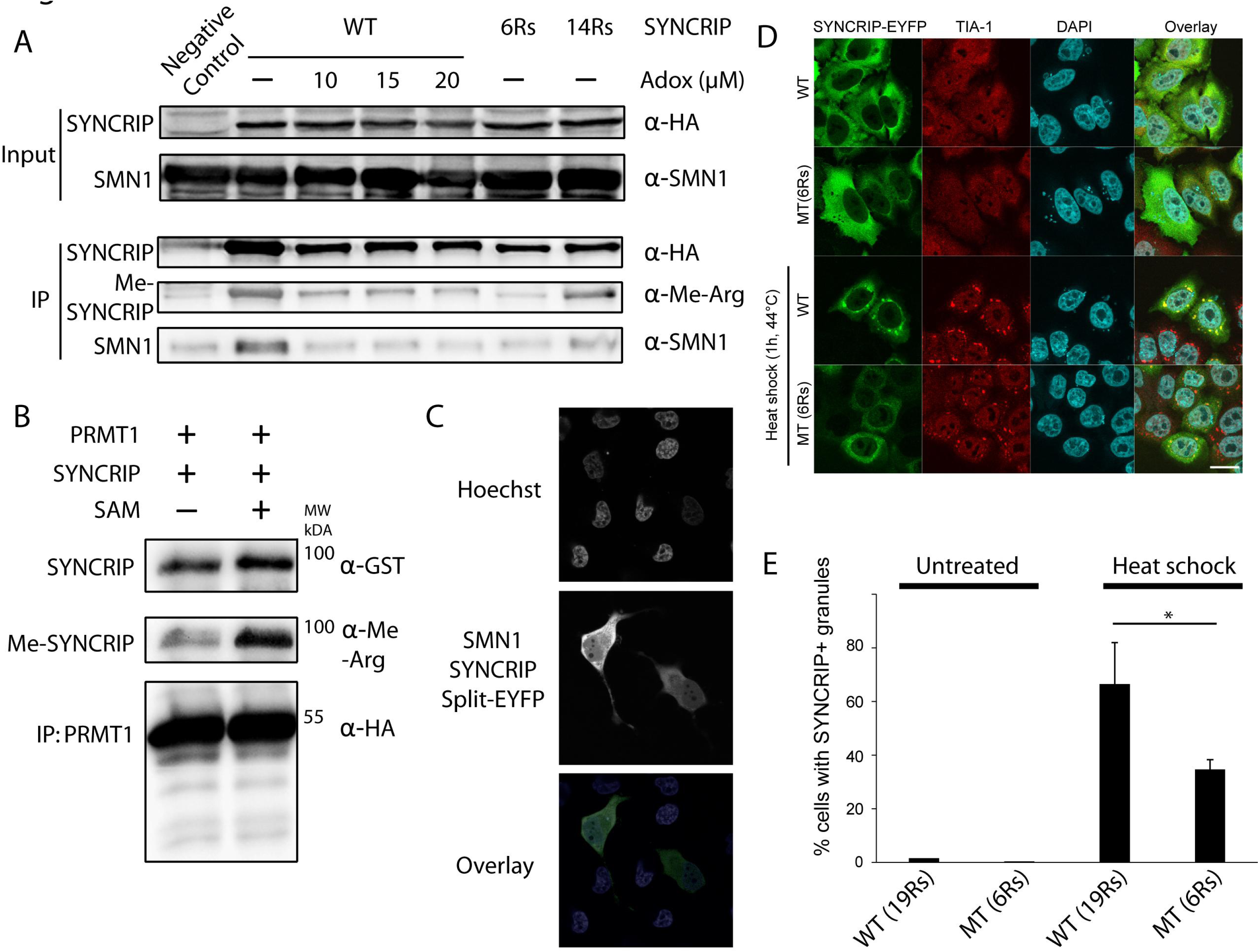
Characterisation of SYNCRIP-SMN1 methylation dependent interaction. A Methylation dependency of SMN1-SYNCRIP interaction. lmmunoprecipitation (IP) of SYNCRIP-HA from HEK293T cells expressing either WT SYNCRIP, or SYNCRIP containing a reduced number of arginines. B SYNCRIP binding and subsequent in vitro methylation by HEK293T cell produced PRMT1 in the presence of co-factor SAM. C Confocal live micrographs of HeLa cells expressing SYNCRIP and SMN1 each tagged with a part of the EYFP fluorophore, Hoescht used to visualize the nucleus. EYFP reconstitutes upon SYNCRIP-SMN1 binding, allowing subsequent visualization. Scale bar indicates 10µm. D Under basal conditions, wild type SYNCRIP and a mutant containing only 6 arginines show the same general sub-cellular localization pattern. Upon heat shock, stress granules (stained with TIA-1 stress granule protein marker) form in both wild type and mutant expressing cells. Wild type SYNCRIP is recruited to stress granules significantly more efficiently than the six arginine SYNCRIP mutant. Scale bar, 20 µm. E Quantification of wild and mutant SYNCRIP recruitment to stress granules. Number of cells with and without SYNCRIP-positive stress granules was counted (n = 100) in four independent experiments. Graph shows mean of four independent experiments, error bars indicate standard deviation. Two sided Mann-Whitney-Wilcoxon Test used to determine statistical significance.*= P<0.05

PRMT1 has been previously shown to bind and methylate SYNCRIP in this C terminal region *in vitro*, making it a strong candidate to mediate the methylation observed here. PRMT1 knock down is toxic to cells and can cause substrate scavenging by other PRMTs, leading to complications in obtaining and interpreting results from standard genetic approaches (Dhar et al. 2013). To ascertain whether PRMT1 produced in live cells is active against SYNCRIP, we purified PRMT1 produced in HEK293T cells for use in an *in vitro* methylation assay. Bacterially produced, and as such highly likely unmethylated SYNCRIP was incubated with PRMT1 immunoprecipitated from HEK293T cells using a STREP-HA tag. As can be seen in Figure 3B, PRMT1 could bind to bacterially expressed, unmethylated SYNCRIP independently of exogenous S-Adenosyl-l-methionine (SAM), the substrate required for methylation. This suggests that neither the co-factor nor priming methylation events are absolutely required for PRMT1 binding. In the presence of SAM the methyl-arginine signal greatly increased indicating SYNCRIP methylation by PRMT1 produced from live cells (Figure 3B).

To characterize the SMN1-SYNCRIP interaction further, we used the split-EYFP system to assay this binding in live cells. SMN1 and SYNCRIP were tagged with N and C terminal sections of EYFP that do not individually fluoresce and co-transfected into HeLa cells. In this system, upon SYNCRIP and SMN1 binding the fluorophore fully reconstitutes allowing direct visualization of the interaction’s sub-cellular localization using live cell imaging. Here we observed that SMN1 and SYNCRIP interact both in the cytoplasm and nucleus (Figure 3C), with a slightly stronger fluorescence in the cytoplasm that is in broad agreement with the localization of each protein when expressed alone or in combination (Supplementary Figure 3). Together these experiments provide further evidence that SYNCRIP is methylated under basal conditions in HEK293T cells and that this methylation leads to direct binding of SYNCRIP to SMN1.

We next moved to investigate the functional relevance of methylated SYNCRIP in human cells. Multiple hnRNPs and SMN1 have been shown to play roles in stress granules formation in cell culture (Guil et al. 2006; Zou et al. 2011). Furthermore, arginine methylation itself has been implicated in stress granule biology, however whether it is a driving force for granule formation or more a function of fully formed stress granules remains unclear (Xie and Denman 2011). We therefore looked to ascertain the sub-cellular localization of wild type and mutant SYNCRIP and endogenous SMN1 under stress and non-stress conditions. We observed stress granule formation under heat shock when expressing both the wild type and mutant SYNCRIP constructs using the endogenous TIA-1 stress granule marker protein (Figure 3D). While wild type SYNCRIP was efficiently recruited to stress granules upon heat shock, the SYNCRIP mutant containing only 6 arginines was only poorly recruited (Figure 3E). Given SYNCRIP is methylated under non-stress conditions, these experiments suggest that arginine methylation is a prerequisite for efficient hnRNP recruitment to stress granules, not a function of the granule stress response. While we could observe overexpressed SMN1 recruitment to stress granules, we could not however observe endogenous recruitment under several stress conditions (Supplementary Figure 4).

### Detailed dissection of methylated and unmethylated arginine array in SYNCRIP

Having validated the importance of the SYNCRIP methyl-R array for SMN1 interaction in cells, we sought to systematically dissect the binding mechanisms of the entire disordered region in both its unmodified and modified states in full. To do so we leveraged the two distinct arms of the arginine methylation regulatory machinery described above, with the methyl-transferase PRMT1 and the methyl-binding protein SMN1 acting as functional readouts for the unmodified and modified disordered regions respectively. Utilizing these two proteins as in-cell molecular probes in the quantitative LUMIER assay would then allow systematic dissection of binding mechanisms of this low structural complexity region.

As the permutations of 19 arginines to lysine mutations is unfeasible to address experimentally (2^19^, >500,000 for position defined permutation), we sought to rationally design mutants based on the cluster proximity of the RG repeats within the array (Supplementary Figure 5). Using site directed mutagenesis we generated a total of 37 mutants in the context of the full length protein that can be designated into three general sub-groups: The first group contains a single, continual stretch of the wild type arginine residues, but the number of arginines and the position of the continual stretch varies across the entire tail (top panel, Figure 4A). Conversely, the second group contains a single, continual stretch of arginine to lysine mutants, but the number and position of lysine mutants in the C-terminal tail is varied (middle panel, Figure 4A). The final smaller group has non-contiguous patches of arginine to lysine mutations distributed across the C-terminal tail (lower panel, Figure 4A).

**Figure 4.**
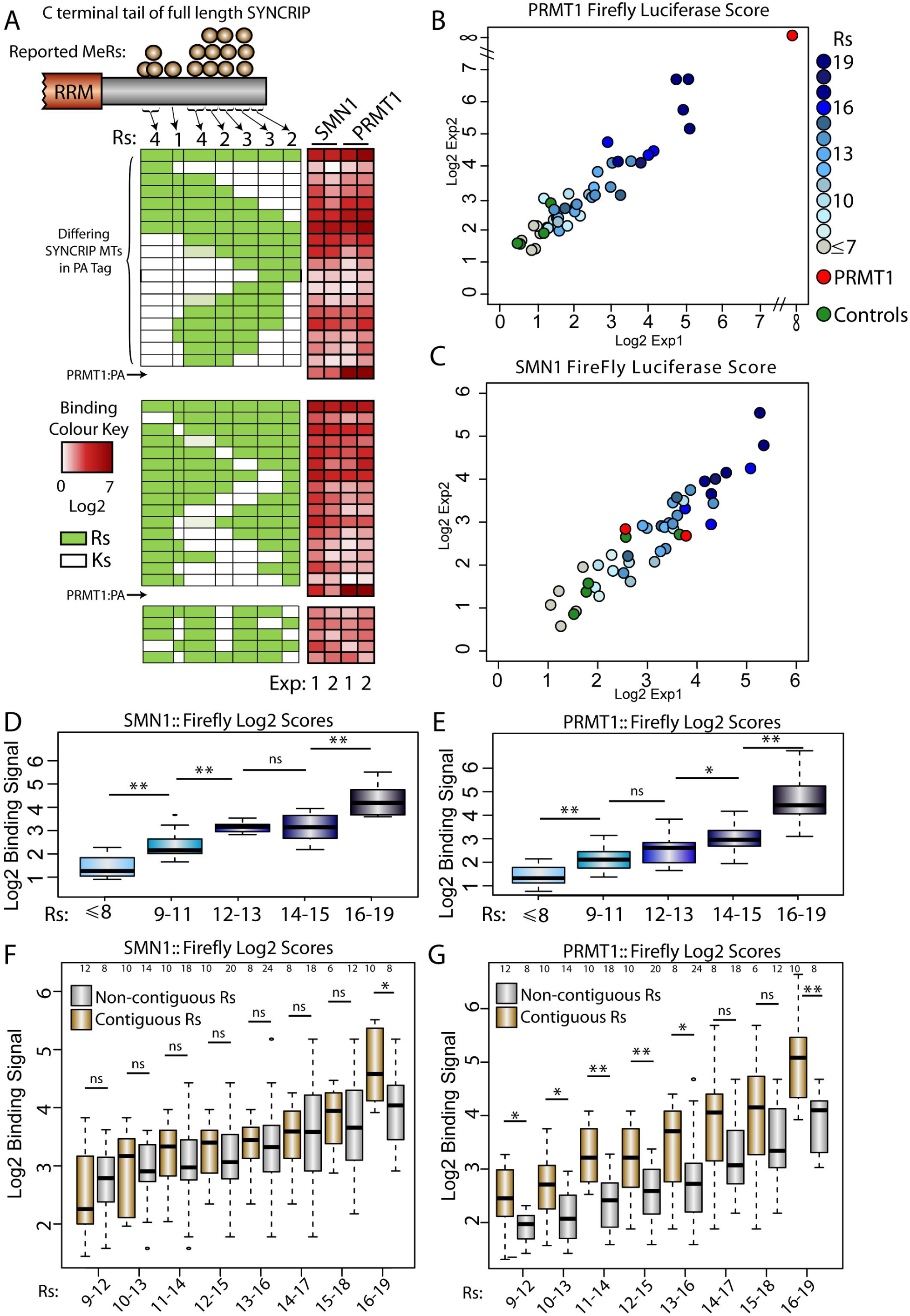
Systematic dissection of arginine requirement for PRTM1 and SMN1 binding to SYNCRIP. A. Schematic diagram representing the R to K full length mutant constructs and their binding signal in each experiment (white-red heatmap next to schematics). Light green boxes represent R patches where only 3 of the 4 arginines were mutated to lysines post sequence verification. PRMT1 (B) or SMN1 (C) binding signals in medium throughput LUMIER experiment assaying multiple R to K mutations on the binding of full length SYNCRIP. SMN1 (D) or PRMT1 (E) binding signal box plots for each group of R to K SYNCRIP mutants. Groups designated by the total number of remaining C terminal arginines. SMN1 (F) or PRMT1 (G) binding signal box plots for sub-groups of R to K SYNCRIP mutants. Each sub-group designated by the total number of arginines in either a contiguous or non-contiguous sequence. Numbers at the top of each blot represents the number of data points in each box plot. One sided Mann-Whitney-Wilcoxon Test used to determine statistical significance. *= P<0.05, **=P<0.01

In agreement with the SYNCRIP mutants used in the endogenous SMN1 immuno-precipitation experiment, the reactivity of a subset of these SYNCRIP mutants with the pan-methyl-arginine antibody correlated well with overall SYNCRIP arginine content. Removing any individual arginine cluster within the array did not abolish methyl-arginine signal, rather the reduction in signal correlated qualitatively with the reduction in methylatable residues (Supplementary Figure 6). As this SYNCRIP arginine to lysine mutant panel can be methylated in a graded manner under basal conditions, it can act as a good proxy for reduced methylation of full length SYNCRIP in cultured cells. We therefore screened each full length mutant for a functional readout of both the unmodified (PRMT1) and modified (SMN1) states of this unstructured region.

Both PRMT1 (Figure 4B) and SMN1 (Figure 4C) LUMIER experiments showed good reproducibility, with mutant expression comparable to wild type SYNCRIP and exhibiting low variability (Supplementary Figure 7). While SMN1 showed only a weak signal for PRMT1 binding that was comparable with controls, PRMT1 show a very strong self-interaction signal, in agreement with previous knowledge on its homo-dimeriseration (Figure 5A and 5B (Zhang and Cheng 2003; Thomas et al. 2010; Weimann et al. 2013)). Through comparing the binding scores with the mutant sequences several trends are immediately clear (Heatmaps next to mutant schematic diagrams, Figure 4A). Mutating either N or C terminal arginines ablated neither SMN1 nor PRMT1 binding completely, only sequentially mutating residues from both terminal groups to leave a small central arginine patch eventually reduced binding to background levels (top panel, Figure 4A). Furthermore, mutation of any individual arginine patch did not reduce the binding signal to background levels with central lysine mutants tolerated in the context of flanking arginines (middle and lower panel, Figure 4A). These experiments suggest a model whereby both the modified and unmodified RG repeat regions mediate their interactions cumulatively, showing an increase binding signal up to restoration of the full 19 wild type arginine residues (grey-blue color code in Figure 4B and C).

**Figure 5.**
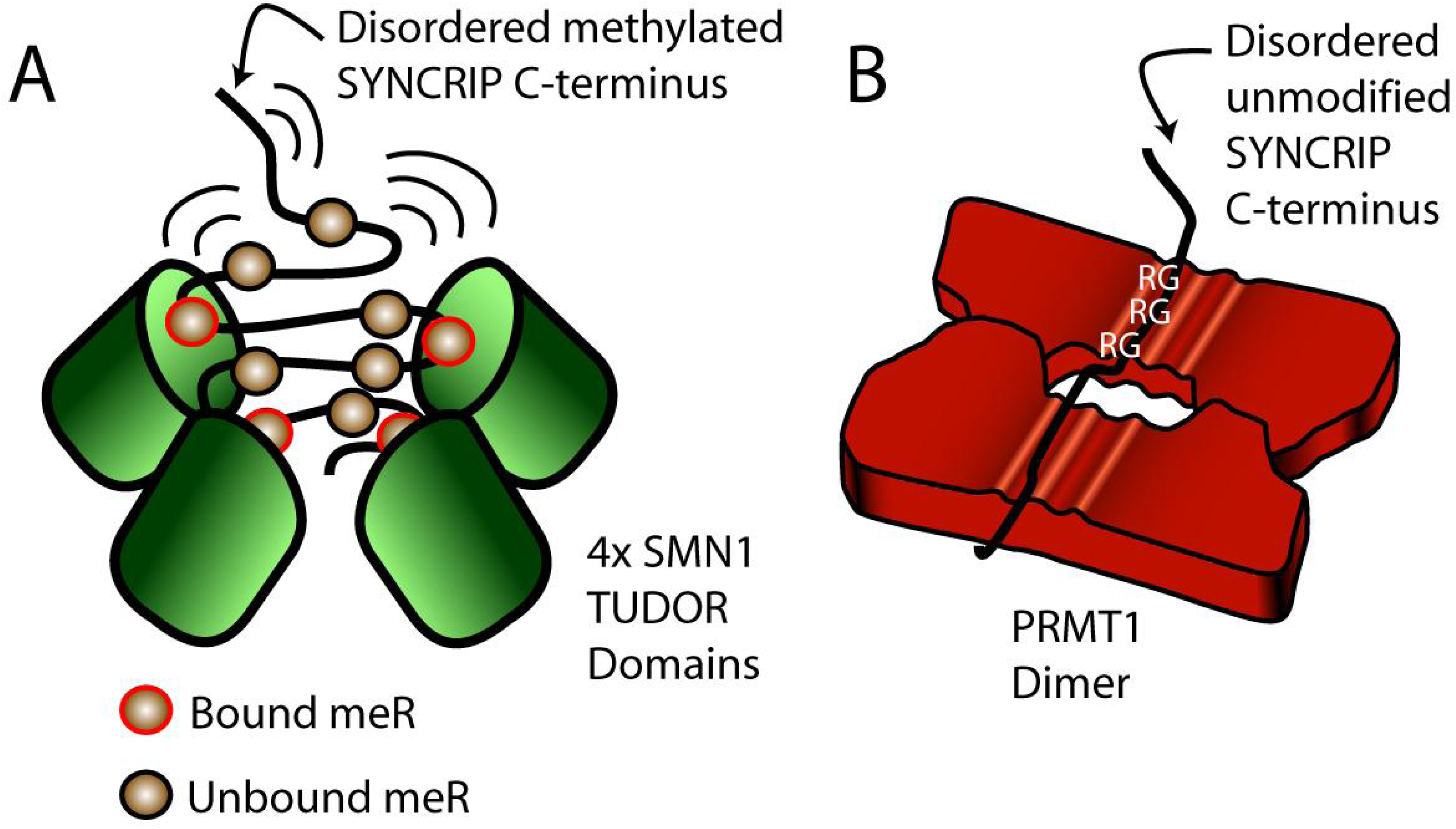
Model of SMN1 methyl-arginine dependent and PRMT1 arginine dependent SYNCRIP interactions. **A** Multiple TUDOR domains in an SMN 1 oligomer independently interact with individual methylated arginine moieties within the flexible SYNCRIP (-terminal tail. B Multiple sequential arginines in the flexible SYNCRIP tail recognized simultaneously by a PRMT1 dimer.

To systematically test whether both modified and unmodified regions follow this overarching model of cumulative arginine dependency we then grouped the mutants based solely on the number of arginines that remain in the C-terminus. In good agreement with this model there is a strong positive correlation between binding score and number of arginines for both SMN1 and PRMT1 (Figures 4D and E). To dissect this further, we then split the SYNCRIP mutants into two sub-categories; one group in which all arginines were present in a contiguous linear sequence, and a second where lysine mutants interrupted the sequence of remaining arginines (non-contiguous). To cover the full spectrum of mutant sub-groups, overlapping levels of total arginine were used to further divide each category. Interestingly, while methyl-arginine reader SMN1 shows little difference between the two groups (Figure 4F), PRMT1 binding signal indicates a clear preference for contiguous arginine stretches, irrespective of the total arginine content (Figure 4G). Importantly, this model refinement still falls within the general cumulative arginine mechanism, as mutants with 15 to 18 arginines present in non-contiguous mutants still show higher PRMT1 binding than mutants with 10-13 contiguous arginine residues (p=0.02, One Sided Mann-Whitney-Wilcoxon test, Figure 4G). Therefore, through leveraging the arginine methylation machinery as in-cell molecular probes we can develop an overall model of cumulative methyl-binding across more than 100 amino acids of disordered protein sequence, and furthermore for the first time differentiate overarching binding preferences of the unmodified and modified RG repeats.

Disordered protein sequences inherently contain little protein structural information and as such are difficult to experimentally investigate. Here we used a functional readout for both the methyl independent (PRMT1) and methylated (SMN1) states to show firstly how a large array of RG repeats can be used to generate a cumulative binding capacity within disordered regions and secondly how PTMs can co-opt these same regions using distinct binding preferences to produce a functional output.

### Discussion

Here we highlighting the dual mechanisms arginine methylation employs to regulate protein function. We identified hundreds of candidates annotated with methyl-R arrays and investigated in detail one of the longest methylated arginine stretches identified, distributed across 19 arginines within the disordered C-terminal tail of the hnRNP SYNCRIP.

As much as 40% of eukaryotic proteomes are annotated as disordered protein sequence (Potenza et al. 2015). Proteins harboring such regions have well established roles in cellular signaling and have been implicated in multiple disease processes (Babu 2016). However, understanding the function of such long unstructured regions in a cellular environment has proved challenging.

To tackle this problem we generated a large set of full length SYNCRIP mutants, allowing investigation into these methyl-R arrays in the context of the full length protein in a quantitative immunoprecipitation assay. This experimental set up does not provide the detailed biophysical data of *in vitro* peptide studies, however the number and length of mutants allows overarching *in vivo* binding principles to be observed that would be otherwise refractory to experimental investigation. In stark contrast to the canonical single PTM-single function paradigm, no individual modified arginine is absolutely required for an interaction in cultured cells. This trend also holds for the modification independent RG repeat functional interaction with PRMT1. Furthermore, the interaction signal observed for modified and unmodified disordered sequences increases as arginines are restored up to the 19 present in wild type SYNCRIP. This finding suggests that the methyl-R arrays identified proteome wide are functionally driven by a requirement for tunable protein interactions.

We could also further refine this cumulative RG motif requirement to be able to propose distinct models for both the modified and unmodified unstructured arrays that relies on the ability of disordered regions to adopt multiple conformations within the cell. Here we show that PRMT1, but not SMN1, substantially prefers contiguous runs of RG motifs within an array. The biophysical properties of PRMT1, SMN1 and their RG peptide recognition mechanisms give clues as to the likely origin of these mechanistic differences. Both SMN1 and PRMT1 are known to oligomerise into higher order structures providing multiple binding sites for each (methylated) arginine in each oligomer (Zhang and Cheng 2003; Burghes and Beattie 2009; Martin et al. 2012). These oligomers are absolutely required for normal functioning and mutants disrupting the SMN1 basic dimer lead to disease phenotypes (Burghes and Beattie 2009). Furthermore, PRMT1 dimerisation is strongly interlinked with AdoMet binding and catalytic activity (Zhou et al. 2015; Thomas et al. 2010), therefore detailed biophysical experiments are required to further disentangle dimerization requirements at each step of the methylation reaction. However, these oligomeric structures allow complex binding mechanisms and provide the basis for the general mechanisms proposed here.

A single SMN1 TUDOR domain monomer can only accommodate binding to one methylated arginine at any given time, yet the arginine binding β-barrel domain still shows an increased affinity for a multi-methylated peptide (Tripsianes et al. 2011). Furthermore, as the TUDOR domain lacks contacts to residues adjacent to the methylated arginine-glycine mark (Tripsianes et al. 2011), repeated methyl-arginine binding can be independent of local sequence context. Therefore, repeated binding does not necessarily require sequential modification along a protein sequence, only methyl-arginines close in 3D space as present here in the long disordered tail region. In the context of multiple TUDOR domains within an SMN1 oligomer, this “one-out one-in” binding mechanism would clearly favor long, multiply modified flexible substrates as each binding pocket could be simultaneously occupied or rapidly rebind dissociated methylated arginines (Figure 5A). In a model with independent recognition events of a single modified residue, multiple binding pockets would aid rapid rebinding of dissociated methylation groups. Furthermore, the interaction would be less sensitive to the linear placement of methylations along a disordered tail that can adopt many conformations, and would consequently mainly be dependent on the total modification level in a confined 3D space. While single methylated arginine residues not present in RG motifs have also been shown to recruit SMN1 (Zhao et al. 2016), we hypothesize that many of the long RG arrays identified here will follow similar cumulative arginine driven binding models.

In contrast to a single methyl-arginine binding to a TUDOR monomer, the PRMT1 monomer contains multiple putative RG binding acidic grooves, three of which have been shown to bind a triple-RGG containing peptide with higher affinity than a single-RGG containing peptide (Zhang and Cheng 2003). This necessarily constrains the arginines in a physically consecutive peptide as multiple motifs across a linear sequence are simultaneously involved in a binding event (Figure 5B). As such, sequence deviation would likely lead to a lower affinity and reduced catalytic activity, in line with previous observations (Zhang and Cheng 2003). All of the long SYNCRIP substrates assayed here contain many multi-RG peptides within a single disordered protein sequence, providing multiple opportunities for PRMT1 oligomer recognition. However, a non-contiguous mutant PRMT1 recognition event is more likely to involve disruptive arginine to lysine flanking mutants than a contiguous mutant with the same arginine content, thus providing a less optimal substrate and the lower binding signals observed here. Given the length of the RG containing arrays identified here, it is plausible that both monomers within a PRMT1 dimer are involved in this recognition event and act in tandem to increase binding strength (Figure 5B).

Long, multiply modified disordered substrates such as the SYNCRIP C-terminal tail fall outside of the classical structure-function paradigm and as such are refractory to direct visualization using standard structural and biochemical approaches. As such, novel approaches are required to untangle exactly how these regions are recognized by the cellular machinery. The genetic approach taken here provides insight into how such long stretches of methyl-arginine residues function within the cell and further how short peptide binding mechanisms translate into overall interaction preferences in the context of a full length protein. These models also provide plausible links to the bioinformatic trends we observed for methyl-R array containing proteins. As protein interactions are driven by protein concentration, tunable interaction mechanisms such as the one described here require tight expression regulation as we observed in the GTEx dataset (Figure 1G). Furthermore, if individual modifications within an array are not absolutely critical for overall function, mutations are less likely to be damaging and subsequently more likely to be observed in population level genetic data (Figure 1H).

This study represents the most comprehensive dissection of extensive low structural complexity regions present proteome wide to date, and furthermore highlight how the cell utilizes two distinct binding mechanisms within these disordered sequences to achieve a similar overarching effect; namely a cumulative contribution of each RG repeat to binding strength.

## Experimental Procedures

### PTM data collation and analysis

Dataset collation was undertaken as in (Woodsmith et al. 2013). Briefly, data for each PTM was obtained from PhosphoSitePlus (Hornbeck et al. 2015) and integrated with publicly available datasets to obtain a non-redundant list of 13 amino acid sequences (13mers). The central amino acid is annotated as modified in each 13mer and only modified lysine or arginine residues were taken forward to the final analysis (Supplementary Table 1).

### Iupred Disorder Analysis

Each RefSeq protein sequence in the analysis was analyzed using the Iupred disorder prediction software (Dosztányi et al. 2005), 0.5 was set as a cut-off to binarise each amino acid into ordered or disordered.

### RBP annotation

RNA binding protein annotation has been shown to be variable depending on the experimental set-up. We therefore took three independent experiment studies (Castello et al. 2012; Baltz et al. 2012; Conrad et al. 2016) for the initial analysis. To annotate the RG array containing proteins as RBPs or not for Figure 2D, we used the list given in (Gerstberger et al. 2014).

### Methyl-R array extraction

>25,000 proteins sequences were computationally scanned in overlapping 20 amino acid windows in the N to C terminus direction. If multiple PTM annotated isoforms were available, the most highly annotated isoform was taken forward. The start of any Methyl-R array was defined as any sequence that contained 3 or more methylated arginines in a 20AA window. The array was then continued unless a 50 amino acid gap between the start of the array and the next methyl-R triplicate appeared. A list of all arrays extracted can be found in Supplementary Table 2.

### Motif Analysis of non-RG driven methylation arrays

We extracted all non-RG methylation sites from arrays driven by methyl-non-RGs (Supplementary Figure 1B). We then used icelogo with default settings (Maddelein et al. 2015) to generate a consensus motif, using as background all methylated arginine sites not present in the positive dataset.

### GTEx dataset

We utilized GTEx_Analysis_v6p_RNA-seq_RNA-SeQCv1.1.8_gene_rpkm.gct for all analyses downloaded from www.gtexportal.org/home/datasets. To aid statistically robustness, we only calculated the median and variance of gene expression for identifiers with >10 samples per tissue, leaving a maximum of 51 tissues per gene identifier. We then used the distribution of the median expression values for the methyl-R array genes as a control for further comparative analysis. We randomly sampled the overall methylation dataset for the same number of genes as present in the methyl-R array target gene set. We then extracted their data from the GTEx dataset, ensuring that for each randomization the distribution of the median expression values across all tissues was statistically indistinguishable from that observed for the methyl-R array genes (Example of distribution comparison Figure 1F left panel). We then compared the distribution of the gene expression variance from the same random sample (Figure 1F, 3 rightmost panels). We repeated the random sampling protocol 100 times, observing the same outcome after each randomization.

### Gnomad dataset analysis

We utilized gnomad.exomes.r2.0.1.sites.vcf.gz for all analyses downloaded from gnomad.broadinstitute.org/downloads, only taking forward the Gnomad annotated isoform that corresponded to the arginine modified isoform from the PTM dataset collation. As a measure for the likelihood of any mutation occurring at a given arginine, we summed the allele frequencies for all mutations per codon across all identifiers. For any given allele frequency cut-off, we then calculated the ratio of the proportion of mutated non-modified arginines to the proportion of mutated modified arginines. We repeated this analysis for three datasets; all proteins targeted by arginine methylation, all proteins targeted by methyl-R arrays and all proteins targeted by methyl-R arrays driven by arginines-glycine motifs.

### Cell Culture

All cell lines were maintained in a humidified incubator at 37°C with 5% CO2. HEK293T cells were used for all immunoprecipitation experiments. For cellular stress experiments, HeLa cells were used for quantification and HeLa Kyoto cells were used for confocal imaging and were grown in DMEM high glucose GlutaMAX (Invitrogen) supplemented with 10% FCS and 10µg/ml gentamicin. For split-EYFP experiments and localization experiments HeLa cells were grown in DMEM containing 10% FCS.

### LUMIER-type experiments

MeBP and SYNCRIP ORFs were transferred to either firefly luciferase-V5 fusion vectors (pcDNA3.1V5-Fire) or protein-A fusion vectors (pcDNA3.1PA-D57), using standard Gateway cloning procedures. For co-IP assays, 3 × 10^4^ HEK293 cells were transiently cotransfected with firefly (75ng) and protein A (PA; 75ng) plasmid DNA using Lipofectamine 2000 (Invitrogen) in each well of a 96-well plate. Cells were lysed 36h after transfection in 100µl HEPES buffer (50 mM HEPES pH 7.4, 150 mM NaCl, 1 mM EDTA, 10% glycerol, 1% Triton X-100 and protease inhibitor (Roche, 11051600)) for 30 min at 4 °C. Protein complexes were precipitated from 80µl cleared cell extract in IgG-coated microtiter plates for 2h at 4°C and rapidly washed three times with 100µl ice-cold PBS. The binding of the firefly-V5–tagged fusion protein (co-IP) to the PA-tagged fusion protein was assessed by measuring the firefly luciferase activity in a luminescence plate reader (Beckmann D TX800, Bright-Glo Luciferase Assay (Promega)). Assays were performed as triplicate transfections. For small scale LUMIER experiments the raw output intensities are displayed for each triplicate. For the methyl-binding protein experiment (Figure 3) the background for SYNCRIP-FIREFLY was calculated as an average of the three lowest reported luminescence readings, converted to a Log(2) scale. This was then subtracted from each reported methyl-binding protein value to be able to observe PA clones that reported robust signals above the background distribution. Two PA proteins (CBX1 and BPRF1) were found to be “sticky” in this experimental set up and were subsequently excluded from further analysis (i.e they showed interactions with a large amount of un-related proteins, data not shown). For the large mutant SYNCRIP experiments, a triplicate of PA untransfected wells were used to estimate the background Log(2) signal in each plate. This was then subtracted from each mutant output value and the Log(2) signals plotted. Furthermore, we checked the observed interaction distribution could not be explained by a simple linear regression of input against output values (R-squared values SMN1-FIREFLY= 0.0196, PRMT1-FIREFLY= 0.0488). Non-interacting controls used to indicate background binding in the large mutant SYNCRIP experiment were U2AF1, BAT3 and SPATA24.

### Stable cell line generation

SYNCRIP and PRMT1 tagged constructs were generated in the pcDNA5/FRT/TO/HA STREP vector using standard gateway cloning (Invitrogen) and transfected into HEK293 cells cultured in DMEM + fetal calf serum (FCS). 48 hours after transfection, transformed cells were selected through incubation with 50µg/ml hygromycin for 12-20 days. Individual colonies were picked and tested for equivalent protein expression induced with 1µg/ml doxycyclin for 24hrs, prior to pooling.

### Endogenous SMN1 immunoprecipitation experiments

For each individual immunoprecipitation, 2.5x10^6^ stable HEK293 cells were seeded in DMEM +FCS (1µg/ml doxycyclin). Each dish was then incubated with the required concentration of Adox or DMSO vehicle control for 24-36 hours. Cells were then lysed in HEPES buffer (as above for LUMIER type experiments), and incubated with pre-blocked anti-HA beads (1% BSA,overnight at 4°C) prior to 3x washing in ice cold lysis buffer. Beads were then re-suspended in 1.5x sample buffer (18 mM Tris-Cl pH 6.8, 0.6% SDS, 3% glycerol, 1.5% β-mercaptoethanol, 0.003% bromophenol blue) prior to electrophoresis and western blot analysis.

### *In vitro* SYNCRIP production

GST tagged SYNCRIP was expressed in 12.5 ml OverNight Express Autoinduction TB-Medium (+Amp, +CAM) at 37°C (150 rpm) for 20h. The bacteria culture was then centrifuged at 1800g (4°C) to collect the cell pellet. The pellet was then re-suspended in 1.85 ml lysis buffer (50mM HEPES, 150mM NaCl, 5% glycerol, 1mM EDTA, 0.5% Brij 58, 1mg/ml lysozyme, 2mM DTT) and incubated on ice for 30 minutes. 350µl Bezonase solution (20mM HEPES, pH8.0, 2mM MgCl_2_, 0.1U/µl Benzoase) was then added to the lysate before a further 30 minute incubation at 4°C and a final centrifugation step at 15,000g for 30 minutes at 4°C prior to the supernatant being stored on ice until further use.

### PRMT1 beads preparation

6 x 10^6^ HEK293 cells expressing either PRMT1 were collected, washed once in ice cold PBS, then incubated on ice for 30 minutes in 0.5ml lysis buffer (50mM HEPES pH8.0, 150mM NaCl, 10% glycerol, 1% triton-X 100). The lysate was then centrifuged at 15,000g, 30 minutes, 4°C prior to the supernatant being inoculated for 1h with pre-washed Strep-Tactin beads suspension at 4°C. PRMT1 beads were then washed four times in lysis buffer prior to being stored on ice until further use.

### SYNCRIP methylation assay

PRMT1 beads were mixed with SYNCRIP bacterial lysate (2:1 by volume) and inoculated shaking (300rpm) for 2h at 30°C either in the absence or presence of 20mM exogenous SAM. The supernatant was removed and stored on ice until further use, remaining beads were resuspended in sample buffer and heated for 5 minutes at 95°C before storage at -20°C prior to western blot analysis. Methylation was detected using anti-mono and dimethyl Arginine antibody (AbCam, [7E6], ab412, raised against asymmetrical NG/NG-dimethyl arginine).

### Localisation experiments

HeLa cells were transfected with FuGene transfection reagent at 3:1 ratio of DNA:reagent using a standard protocol. Live cell imaging of split-EYFP was undertaken on MatTek dishes 22hrs post transfection with 10 minutes of Hoescht incubation prior to visualization. For individual and co-localisation experiments, cells seeded on glass coverslips were fixed 16 hours post transfection with 4% paraformaldehyde. EYFP signal was detected using a chicken anti-GFP antibody (abcam, ab13970) followed by an anti-chicken Alexa-Flour-488 secondary (Thermo). PA signal was detected using rabbit IgG (Santa Cruz) followed by anti-rabbit Alexa-Flour-555 secondary (Thermo).

### Confocal microscopy and image processing for cellular localisation experiments

Confocal laser scanning microscopy was performed on a Fluoview 1000 confocal microscope (Olympus) equipped with a UPLSAPO60/1.3 numerical aperture silicon immersion oil immersion lens. Images were taken with the following excitation (Ex) and emission (Em) settings: Hoechst Ex: 405 nm diode laser (50mW) Em: 425–475 nm; GFP, AlexaFluor488 Ex: Multi-Line Argon laser 488nm (40mW) Em: 500–545 nm; EYFP, Ex: Multi-Line Argon laser 515 nm (40mW) Em: 530–545 nm; AlexaFluor555 Ex: 559nm diode laser (20mW) Em: 570–625 nm.

### Cellular Stress Experiments

In order to avoid formation of stress granules by overexpression, low amounts (10 ng per 24-well) of DNA coding for EYFP-SYNCRIP (WT/6R) was cotransfected with an empty vector (pcDNA3.1-hygro(+); 490 ng per 24-well) using Turbofect (Fermentas) according to the manufacturer’s instructions. Stress treatment was carried out either by heat shock (1 h at 44°C) or by addition of 0.5 mM sodium arsenite (30 min at 37°C). Cells were immediately fixed in 3.7% formaldehyde in PBS for 7-10 min and permeabilised with 0.5% Triton X-100 in PBS. Cells were blocked for 10 min in blocking buffer (1% donkey serum in PBS/0.1% Tween-20) and incubated with primary antibody (Monoclonal mouse anti GFP (for detection of YFP-SYNCRIP): Hybridoma was kindly provided by A. Noegel, Cologne, Germany (Noegel et al. 2004); purified antibody was gift from M. Kiebler, LMU, Munich. Mouse anti SMN: BD (610646), goat anti ia-1 (G3), Santa Cruz (sc-166247)) and secondary antibodies (Invitrogen/Molecular Probes) diluted in blocking buffer. Washing steps were performed with PBS/0.1% Tween-20. Nuclei were stained with DAPI (0.5 µg/ml, Sigma) and mounted in prolong diamond mounting medium (Invitrogen) before analysis by confocal fluorescence microscopy.

### Confocal microscopy and image processing for cellular stress experiments

Images for colocalisation of YFP-tagged SYNCRIP with stress granule markers were acquired by confocal microscopy on an inverted Leica SP8 microscope (Bioimaging core facility of the Biomedical Center), equipped with lasers for 405, 488, 552 and 638 nm excitation. Images were acquired with a 63x1.4 oil objective, image pixel size was 59 nm. The following fluorescence settings were used for detection: DAPI: 419-442 nm, Alexa488/YFP: 498-533 nm, Alexa555: 598-634; Alexa647: 650-700 (for quadruple stain) or 649-698 (for triple stain). Recording was sequentially to avoid bleed-through using a conventional photomultiplier tube. Confocal images were acquired using LAS X (Leica) and processed using Image J software applying linear enhancement for brightness and contrast. For illustration of the localization of SMN in response to arsenite treatment, a stack of 10 (unstressed) or 12 (+arsenite) sections in 300 nm step size was acquired and projected using the maximum intensity projection function in the LAS X software.

## Author Contributions

US supervised the project. JW conceived the project. JW undertook the bioinformatic analysis, SYNCRIP mutagenesis and all LUMIER-type experiments. VT and CH provided reagents. VC and NB generated stable HEK293 cell lines, VC performed the endogenous SMN1 immunoprecipitation, NB performed the *in vitro* methylation. RE and SH/CAA performed the microscopy under the supervision of OR and DD respectively. JW and US wrote the manuscript with input from all co-authors, JW generated the figures.

---

**Supplementary Figure 1.**
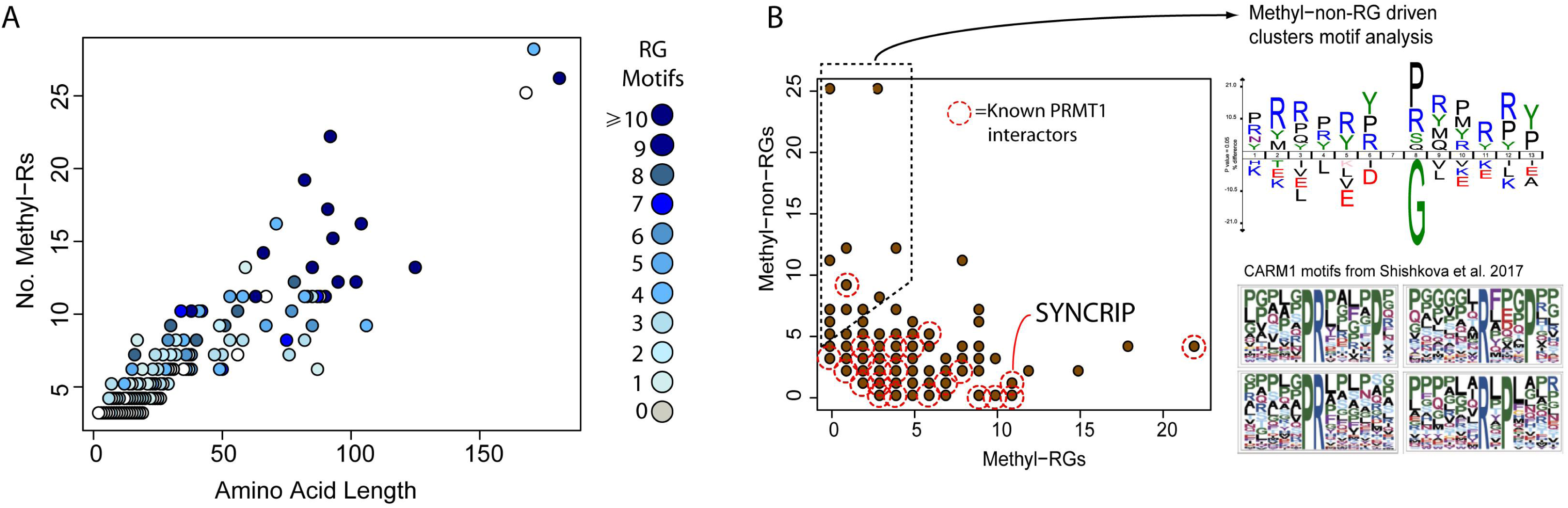
Methyl-R Window General Characteristics and detailed depiction SYNCRIP (-terminus. A Scatterplot depicting the total length in amino acids and number of methyl-Rs contained in the identified clusters of arginine methylation. B Scatterplot depicting the distribution of non-RG and RG methylation sites in the identified clusters of arginine methylation. Each brown point can represent several proteins. Right of plot; sequence motif of all non-RG sites in methylation clusters indicated by dashed box for comparison with published CARM1 motifs.

**Supplementary Figure 2.**
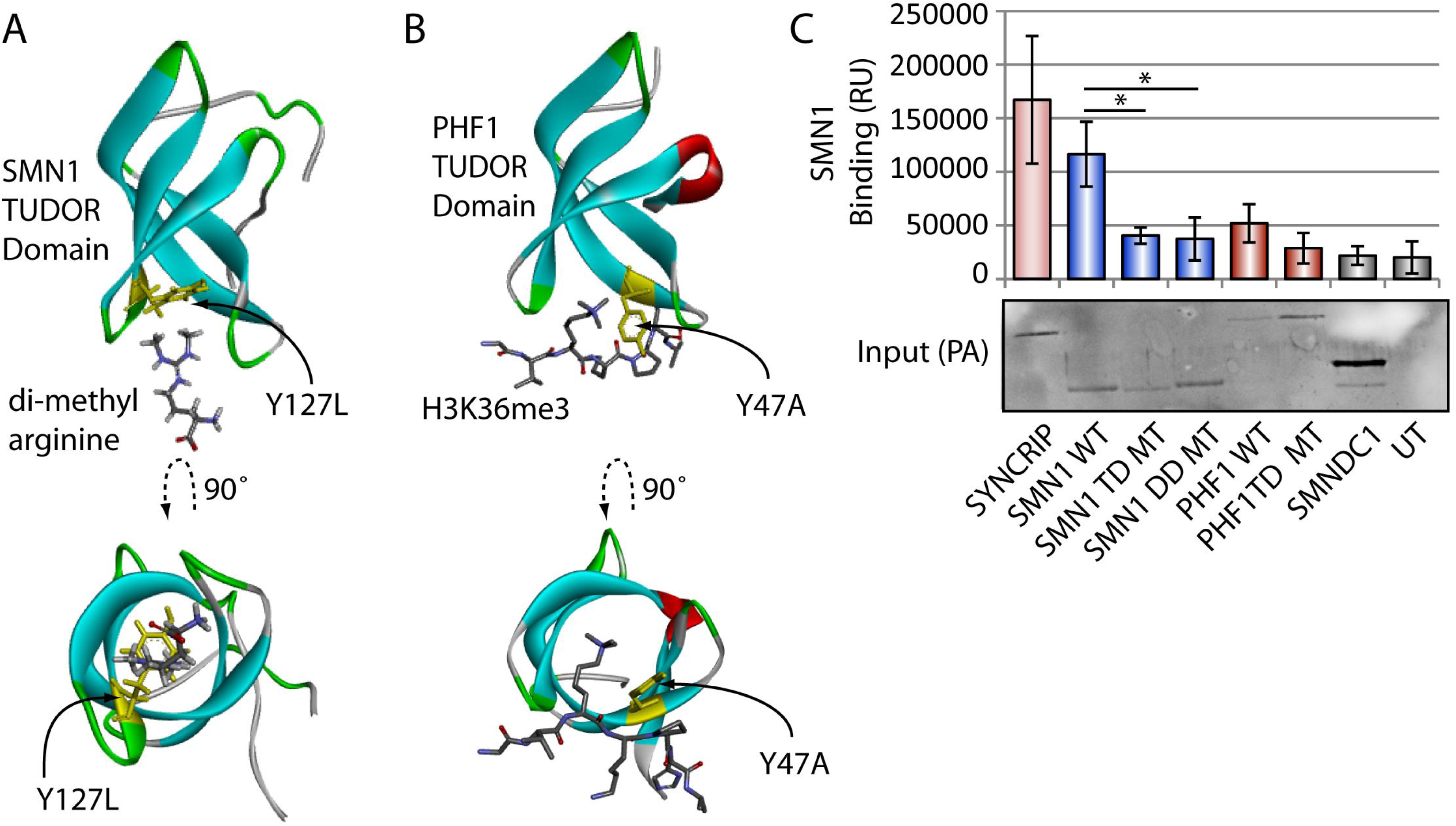
**A** Structure of SMN1 TUDOR domain monomer and binding mutant used in this study (Tripsianes, K. NSMB, 2011, PDB 4AEA). **B** Structure of PHFTUDOR domain and binding mutant (Musselman CA, NSMB, 2012, PDB 4HCZ). **C** SMN1 dimerisation LUMIER experiment. LUMIER experiment testing SMN1 and PHF1 mutants against wild type SMN1. TD = TUDOR domain, DD = Dimerisation domain. Experiments undertaken in triplicate, with error bars representing highest and lowest observed values. Mann-Whitney-Wilcoxon Test used to determine statistical significance.*= P<0.05

**Supplementary Figure 3.**
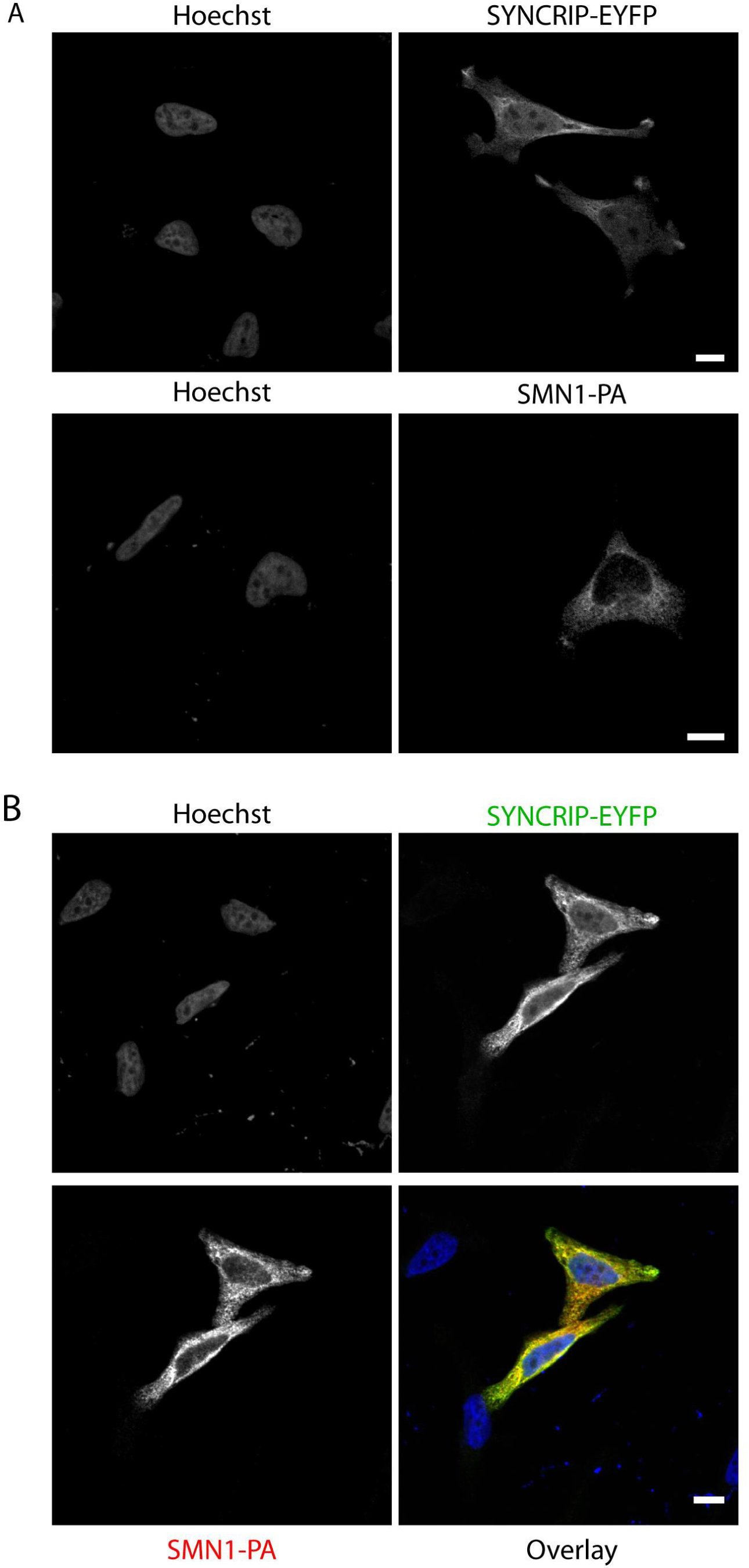
SMN1-PA and SYNRIP-EYFP distribute diffusely in the cytoplasm and nucleus. A Hela cells transfected individually with either SYNCRIP-EYFP or SMN1-PA were fixed and subjected to immunofluorescence microscopy using antibodies against GFP or protein-A. B Hela cells transfected simultaneously with SYNCRIP-EYFP and SMN1-PA were fixed and subjected to immunofluorescence microscopy using antibodies against EYFP or protein A. Hoescht was used to stain the nucleus. Scale bar indicates 1Oµm.

**Supplementary Figure 4.**
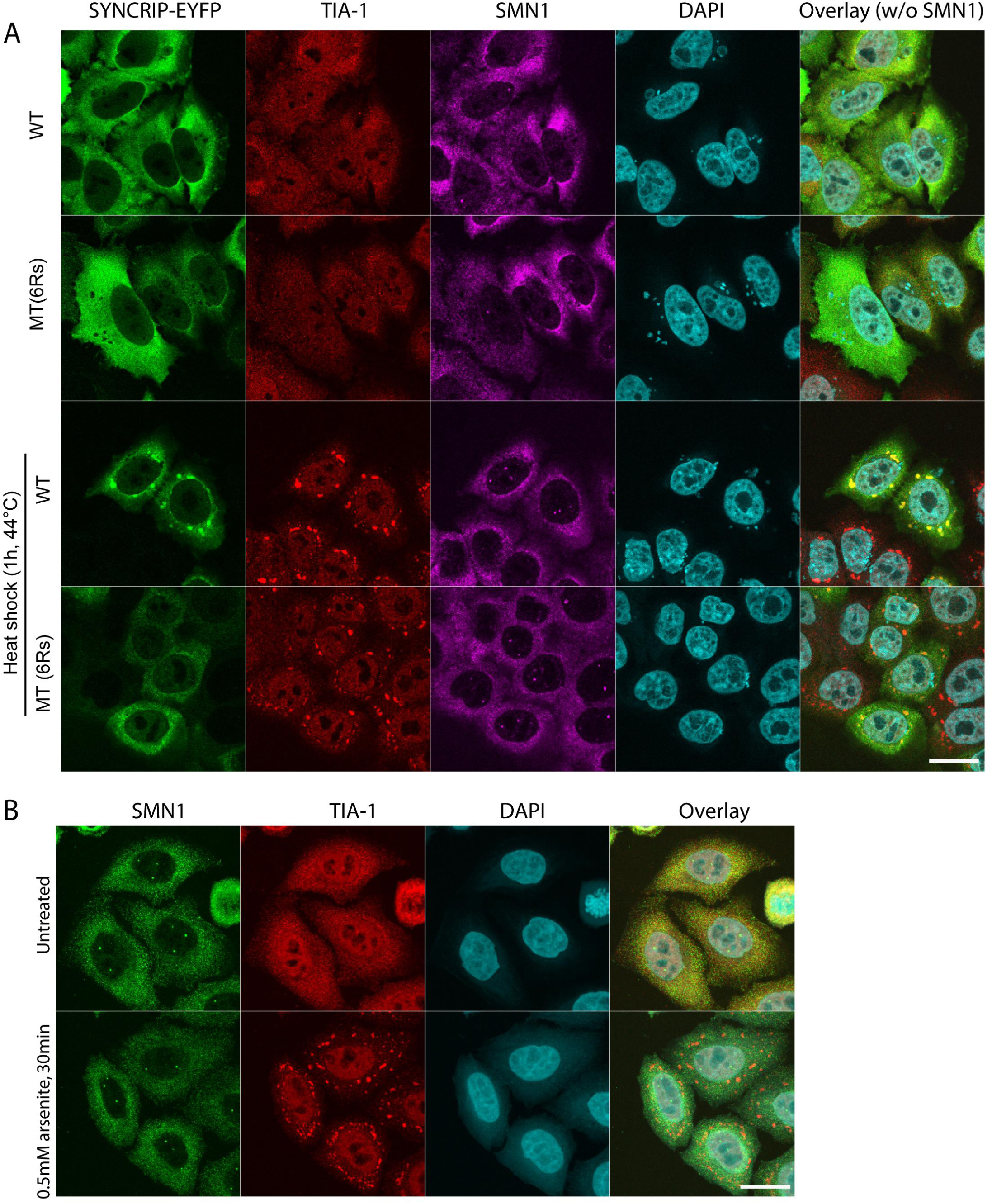
Endogenous SMN1 is not recruited to stress granules in HeLa cells after diverse stresses. A Extended panel from Figure 3D. He La cells, either untreated or incubated for 1 h at 44^°^C were stained for EYFP (using a GFP antibody), endogenous SMN and TIA-1 as marker for stress granules and analysed by confocal microscopy. B Hela cells, either untreated or incubated for 30min with 0.SmM arsenite were stained for endogenous SMN and TIA-1 as marker for stress granules and analysed by confocal microscopy. DAPI used to stain the nucleus. Scale bar indicates 20 µm.

**Supplementary Figure 5.**
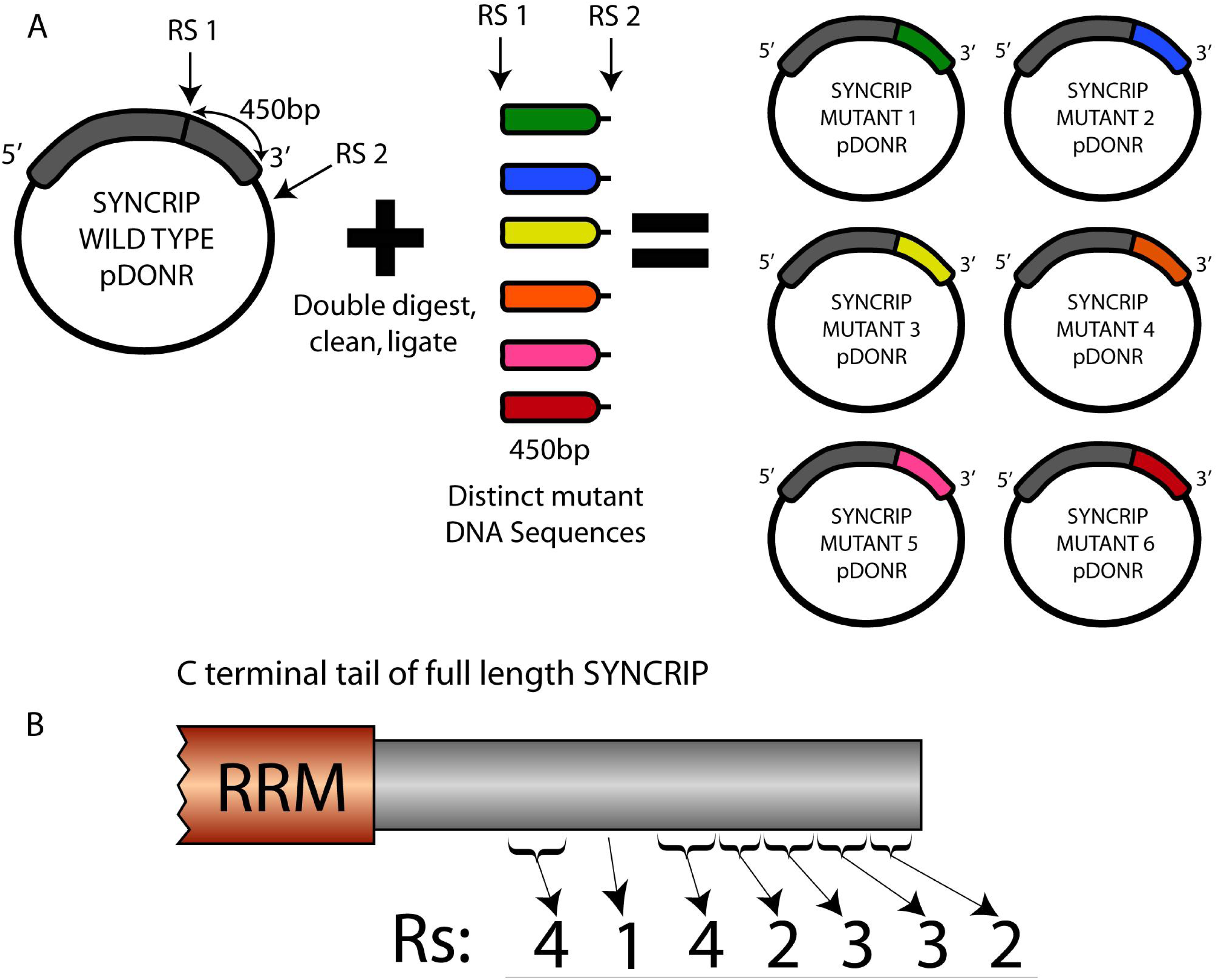
SYNCRIP mutant generation and initial SYNCRIP mutant LUMIER experiment. A Schematic representation of full length SYNCRIP mutant generation. The C terminal tail of SYNCRIP was digested out of the pDONR vector using one internal and one external restriction site (RS1 and RS2). This vector was gel purified and then ligated with individually synthesized DNA fragments to generate full length, in frame SYNCRIP mutants. B Number of arginine residues in each designated mini-cluster in the C terminal tail of SYNCRIP

**Supplementry figure 6.**
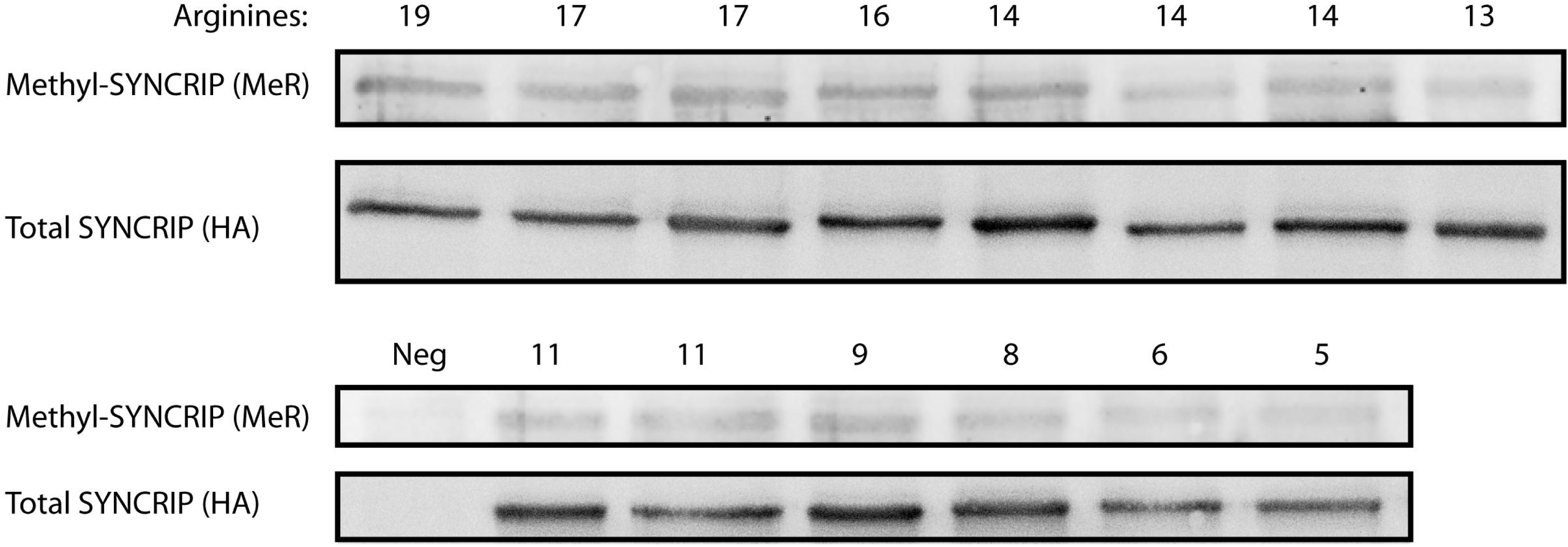
Quantification of SYNCRIP mutant methylation levels. Precipitated HA-STREP tagged SYNCRIP MTs blotted for total levels and methylated arginine levels.

**Supplementary Figure 7.**
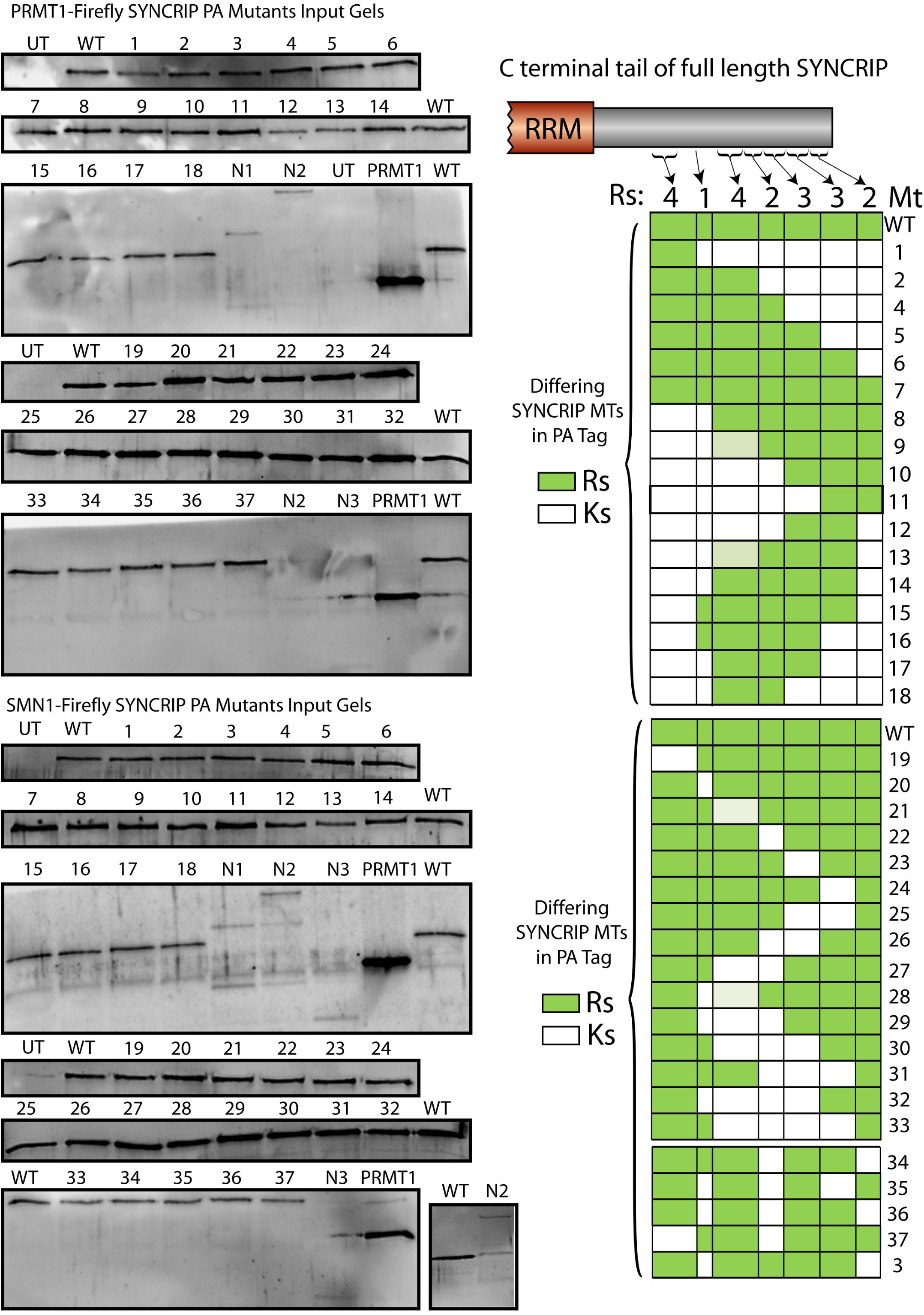
Total protein levels of SYNCRIP mutants. Each SYNCRIP MT in the large LUMIER-type experiment blotted for total protein levels using an anti-protein A antibody.

